# Intricate interactions between antiviral immunity and transposable element control in *Drosophila*

**DOI:** 10.1101/2024.03.18.585529

**Authors:** Camille A Mayeux, Anaïs Larue, Daniel S. Oliveira, Marion Varoqui, Hélène Henri, Rita Rebollo, Natacha Kremer, Séverine Chambeyron, Marie Fablet

**Affiliations:** Laboratoire de Biométrie et Biologie Evolutive; Université Lyon 1; CNRS; VAS; UMR 5558, Villeurbanne, France; Univ Lyon, INRAE, INSA-Lyon, BF2I, UMR 203, 69621 Villeurbanne, France; Institute of Biosciences, Humanities and Exact Sciences, São Paulo State University (Unesp), São José do Rio Preto, São Paulo, 15054-000, Brazil; Institute of Human Genetics, UMR9002, CNRS and Université de Montpellier, Montpellier, France; Institut Universitaire de France

## Abstract

Transposable elements (TEs) are parasite DNA sequences that are controlled by RNA interference pathways in many organisms. In insects, antiviral immunity is also achieved by the action of small RNAs. In the present study, we analyzed the impacts of an infection with Drosophila C Virus (DCV) and found that TEs are involved in a dual response: on the one hand TE control is released upon DCV infection, and on the other hand TE transcripts help the host reduce viral replication. This discovery highlights the intricate interactions in the arms race between host, genomic parasites, and viral pathogens.

**Significance statement:** Transposable elements (TEs) are widespread components of all genomes. They were long considered as mere DNA parasites but are now acknowledged as major sources of genetic diversity and phenotypic innovations. Using *Drosophila* C virus, here we show that TEs are at the center of defense and counter-attack between host and virus. On the one hand, TE control is released upon viral infection, and on the other hand, TE transcripts help the host reduce viral replication. To our knowledge, this is the first time such a complex host-pathogen interaction involving TEs is shown.

## Introduction

Transposable elements (TEs) are DNA sequences considered as genomic parasites because they have the ability to move and multiply along chromosomes at the expense of the host. TEs are very diverse in structure (number of genes, length, presence of repeats, *etc*.) and abundance, from a few percent in the honeybee *Apis mellifera* (1) to ∼ 80% of the recently sequenced arctic krill genome (2). Tolerance to TEs is possible thanks to epigenetic mechanisms that inhibit their activity, and there is a strong selective pressure for these mechanisms to be particularly efficient in gonads, where the genomes of the next generation lie (3–5). In insects, this control is essentially achieved through RNA interference (RNAi) pathways, and in particular *via* the piRNA pathway. piRNAs are 23-30 nt-long single-stranded RNAs that target TE sequences through sequence complementarity. They form complexes with proteins displaying RNAse activity or triggering heterochromatinization (6–8).

RNAi is also the first line of defense against viruses in Arthropods (9–12). Antiviral immunity relies on small RNAs known as siRNAs, which are 21 nt-long single-stranded RNAs, produced by Dicer-2 from double-stranded RNA templates. Whereas siRNAs are known to play a role in the somatic control of TE activity (13, 14), the intricate interplay between TE regulation and antiviral immunity through RNAi remains largely unexplored. Upon viral infection, viral RNA genomes are converted to DNA by TE reverse transcriptases, and this process leads to the increased production of viral siRNAs, facilitating the establishment of persistent infection (15, 16). Moreover, the activation of the *mdg4* TE at the pupal stage was recently found to prime the host’s antiviral immunity for adult stage (17). In addition, in a previous investigation in *Drosophila* using the Sindbis virus (SINV), which naturally infects mosquitoes, we demonstrated that the host antiviral response triggered an amplified production of TE-derived small RNAs. This resulted in the reduction of TE transcript amounts (18). In the present study, we investigated the more closely intertwined interaction between the *Drosophila* host and its natural pathogen Drosophila C virus (DCV).

DCV is a non-enveloped virus that belongs to the Dicistroviridae family. It has a 9.2 kb-long, single-stranded RNA genome of positive polarity (19). DCV is a natural pathogen of *Drosophila* (20, 21), and is horizontally transferred through oral infection. A few days after infection, and independently of the route of infection –either oral or systemic through experimental injections–, DCV viral particles can be detected in the fat body, trachea and visceral muscle of the crop, midgut, hindgut, and gonads (22). Systemic DCV infection causes intestinal obstruction that leads to fly death (23). DCV was first described in *D. melanogaster* but it can also naturally infect *Drosophila simulans* (20). The 99 most N-terminal residues of DCV ORF1 –encoding non-structural proteins– correspond to 1A, a viral suppressor of RNA interference (VSR) that binds long double-stranded RNAs (dsRNAs) and therefore prevents Dicer2-mediated production of siRNAs (24, 25).

Here we show that DCV infection disrupts the regulation of TEs in a more intricate way than what was observed with SINV (18). This is particularly pronounced in the *D. melanogaster* strain, as compared to the *D. simulans* strain. In addition, we show that hosts with a higher TE load in the transcriptome display a more efficient antiviral immunity. This study highlights the prominent contribution of TEs in the immune response of *Drosophila*.

## Results and Discussion

### Host response to DCV infection in *D. melanogaster w^1118^* and *D. simulans* Makindu

We infected flies by intra-thoracic pricking (26), and followed fly mortality and DCV replication. The infection rapidly led to fly death, and significantly more rapidly in *D. simulans* Makindu (log-rank test, p-value = 0.008). On average, half of the flies were dead at 5.4 days post infection (dpi) in *D. melanogaster w^1118^,* and 3.8 dpi in *D. simulans* Makindu (Fig. 1A). Accordingly, RT-qPCR revealed that DCV replication was 2.6-fold higher in *D. simulans* Makindu compared to *D. melanogaster w^1118^* at 4 dpi, which corresponds to the peak of viral titer (Fig. 1B).

**Figure 1.**
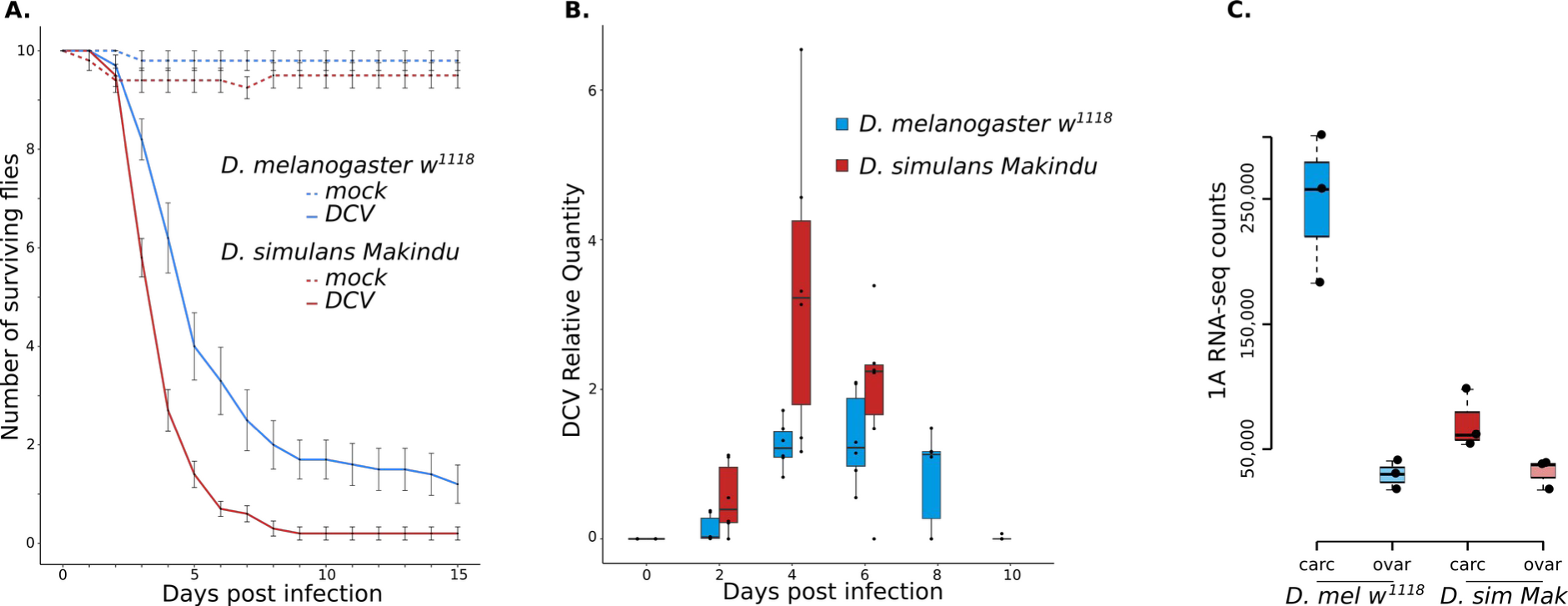
Response to DCV infection in *D. melanogaster w^1118^* and *D. simulans* Makindu. **A.** Fly survival upon DCV infection. **B.** Kinetics of DCV titers followed using RT-qPCR. **C.** Raw RNA-seq read counts mapping against 1A sequence. RNA-seq read mapping along the sequence of DCV genome is shown in Fig S1.

We performed RNA-seq and small RNA-seq experiments at the peak of viral titer, *i.e.,* 4 dpi in both species, in dissected ovaries and the rest of the body –hereafter called “carcasses”. DCV RNAs were highly abundant among the sequenced molecules in carcasses. We could also detect DCV RNAs in ovaries, however in much lower amounts, potentially corresponding to the presence of DCV in the muscle cells of the ovarian peritoneal sheath, as described by Ferreira *et al.* (22).

As previously described (23), DCV infection by pricking leads to many down-regulated genes in *D. melanogaster w^1118^*somatic tissues, as well as in *D. simulans* Makindu. In particular, down-regulated genes were enriched in genes related to metabolic functions, whereas the few activated genes were enriched in stress-response activity, as reported by Chtarbanova *et al.* (23), but also in immune processes (Fig S2).

It should be noticed that we cannot directly draw conclusions on a *D. melanogaster versus D. simulans* species comparison based on only one strain per species. Instead, it would require the inclusion of several strains for each species. Nevertheless, we have to acknowledge the differences in patterns between *D. melanogaster w^1118^* and *D. simulans* Makindu, which could correspond to several hypotheses. i) The sequencing data were produced at 4 dpi for both strain, which corresponds to a higher number of dead flies and a stronger viral replication in *D. simulans* Makindu. Therefore, 4 dpi seems to be further along in the infection process in *D. simulans* Makindu. ii) It is possible that the infection protocol leads to acute infection in *D. simulans* Makindu while persistence is achieved in *D. melanogaster w^1118^*. iii) DCV was first described as a *D. melanogaster* pathogen. The differences between *D. melanogaster* and *D. simulans* outcomes may be due to *D. simulans* not being its preferential host, even though DCV has been found in natural samples of *D. simulans* (20).

### DCV infection impacts the transcript amounts of transposable elements

We analyzed the amounts of transcripts for 237 TE families in the carcasses of the different samples, and we found a global significant increase upon DCV infection in *D. melanogaster w^1118^* (median per TE family log2 Fold Change (log2FC) = 0.28, paired Wilcoxon test: p-value = 8e-12). This is in agreement with our previous reanalysis of data produced from *D. melanogaster y^1^* fat bodies upon DCV infection, which also showed a clear increase in TE transcript amounts (18, 27). In the present data, forty three TE families are significantly upregulated upon DCV infection (DESeq2 adjusted p-value < 0.05), whereas seven are significantly down-regulated (Fig. 2A; See Fig S3 for TE family details). We could not detect significant differences across TE classes (Table S4). In *D. simulans* Makindu, there is virtually no modulation: we observed a significant increase for six TE families and a significant decrease for four TE families (Fig. 2B).

**Figure 2.**
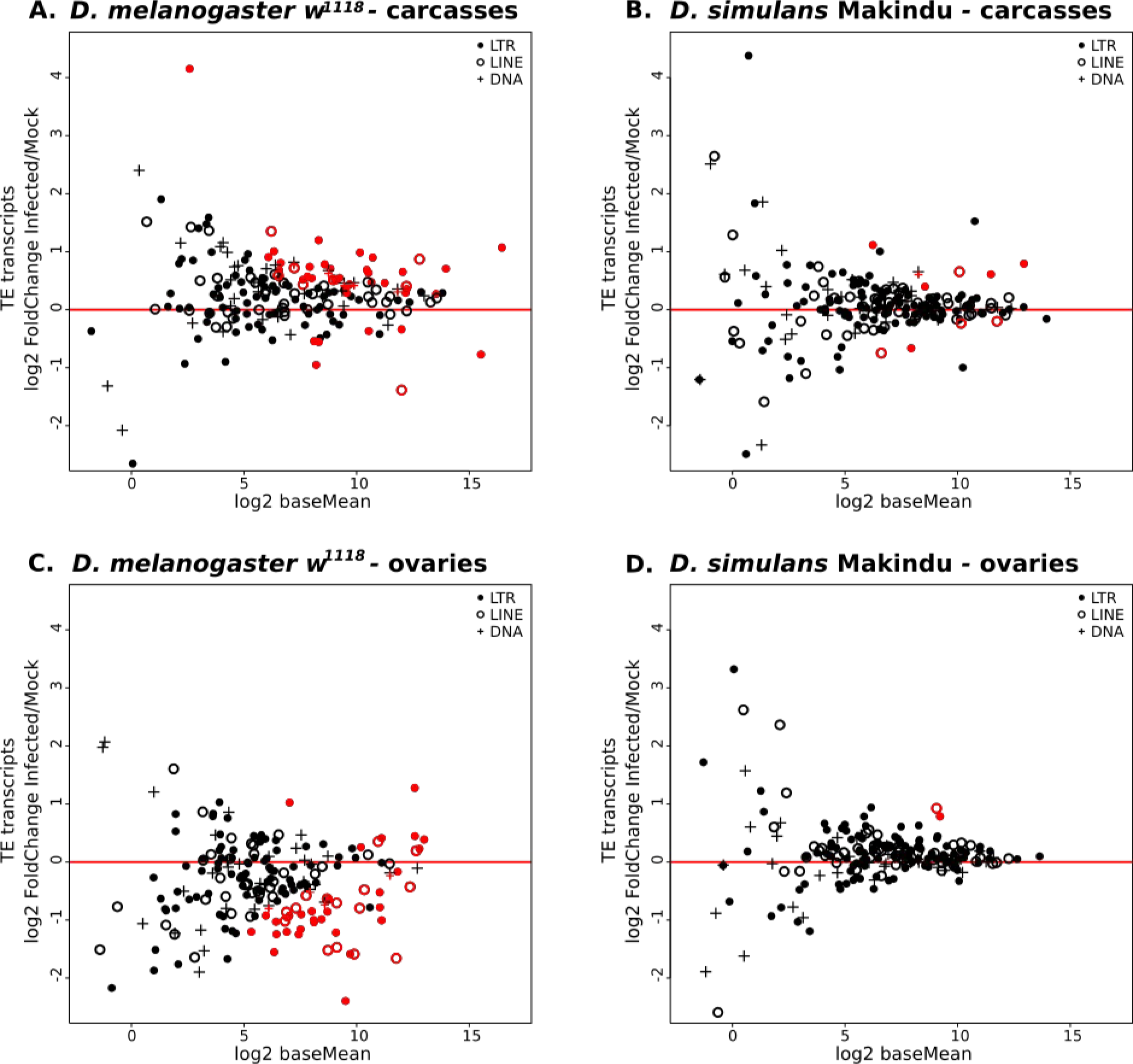
TE transcript modulation upon DCV infection. **A.** TE transcripts log2FC between DCV infected and mock conditions. Dot shapes indicate TE classes, red line is log2FC=0, *i.e.* no modulation. Red dots are TE families displaying significant differential expression at the 0.05 level for DESeq2 adjusted p-values. **A.** *D. melanogaster* w^1118^ carcasses, **B.** *D. simulans* Makindu carcasses, **C.** *D. melanogaster* w^1118^ ovaries, **D.** *D. simulans* Makindu ovaries.

In ovaries, we observed an opposite pattern for *D. melanogaster w^1118^* samples, with a clear decrease of TE transcript amounts: median per TE family log2FC = −0.38 (paired Wilcoxon test : p-value = 1e-9). Transcript amounts increased significantly for nine TE families whereas they significantly decreased for 41 TE families (Fig. 2C). In *D. simulans* Makindu, DCV infection had very little impact on ovarian TE transcript amounts (median per TE family log2FC = 0.10, paired Wilcoxon test : p-value = 5e-9) (Fig. 2D).

In order to understand the mechanisms underlying changes in TE transcript amounts upon DCV infection, we analyzed small RNA-sequencing data from the same experimental conditions. The data showed the expected patterns in size distribution and nucleotide composition (Fig S5-S6). Ping-pong signatures were detected in all conditions, indicating that flies infected with DCV could still produce functional piRNAs (Fig S7). TE-derived siRNAs and piRNAs of both polarities clearly increased upon DCV infection in carcasses of *D. melanogaster w^1118^* and *D. simulans* Makindu, whereas we detected very low size effects in ovaries (Fig S8-S10). The increase was stronger in piRNAs compared to siRNAs (all TE families together, median log2FC: piRNAs: [0.25; 1.28], siRNAs: [0.05, 0.75], Fig S11).

When TE control is achieved *via* small RNA interference, the expectation is that an increase in small RNAs leads to a decrease in RNA amounts, and reciprocally. However, here, the TE families displaying small RNA increase upon infection are globally not significantly enriched in downregulated families, and this was consistent across all conditions (Table S12). Such a result reveals a dissociation between TE transcript amounts and TE-derived small RNAs in the present experimental system.

It has been demonstrated that viral infections in *D. melanogaster* trigger the active uptake of dsRNA molecules of viral origin in haemocytes, which fuels the siRNA machinery and allows the systemic production of antiviral siRNAs (16, 28). Using Sindbis virus (SINV), we previously proposed that dsRNA molecules of TE sequences hitchhike the uptake pathway, and are opportunistically imported within haemocytes along with viral dsRNAs. This would then allow the enhancement of TE-derived small RNA production and explain the observed systemic increase of TE-derived small RNAs, which leads to a decrease in TE transcript amounts (18). Here, we found a global increase of both TE RNAs and TE-derived small RNAs in *D. melanogaster w^1118^* carcasses. We propose that such a discrepancy is due to 1A, the VSR encoded by DCV (no VSR has been described for SINV). In the tissues where DCV is expressed, 1A is produced and binds long dsRNAs, however without any sequence specificity (24, 25, 29). Therefore, 1A mostly binds viral dsRNAs, but also other dsRNAs such as those produced from TE sequences, either due to antisense transcripts annealing to sense RNAs or to single-stranded transcripts folding back on repeated regions. Eventually, the production of TE-derived siRNAs is prevented, resulting in increased TE RNA amount. This is in agreement with the strong increase in TE transcript amounts that we previously observed in *dcr2* mutants upon SINV infection (18). Therefore, we propose that 1A is responsible for the decorrelation between TE RNA and siRNA amounts observed above. TE-derived small RNA amounts would thus result from these two mechanisms with opposed effects: the activation of the dsRNA uptake pathway leads to piRNA and siRNA increase while the action of 1A leads to the reduction of siRNA production. Indeed, since 1A specifically binds dsRNA molecules, it is expected to interfere mainly with siRNA production, which explains the resulting increase in small RNAs is lower for siRNAs compared to piRNAs. In the case of the previously studied SINV infection, the tripartite interaction between host, virus and TEs is simpler and only results from TEs hitchhiking the dsRNA uptake pathway. Here, the production of a VSR adds another layer of interaction. This ends up in an intricate arms race between the virus, the host and its TEs (Fig. 3). Moreover, we propose that this explains the virtual absence of any TE phenotype in *D. simulans* Makindu. Indeed, the RNA-seq data reveal that the number of reads mapping against the sequence corresponding to 1A is the highest in *D. melanogaster w^1118^* carcasses, and very low in ovarian samples. In carcasses, the number of reads mapping against 1A is significantly higher in *D. melanogaster w^1118^* compared to *D. simulans* Makindu (Fig 1C). These species-specific differences may be due to contrasted efficiencies in the production of 1A.

**Figure 3.**
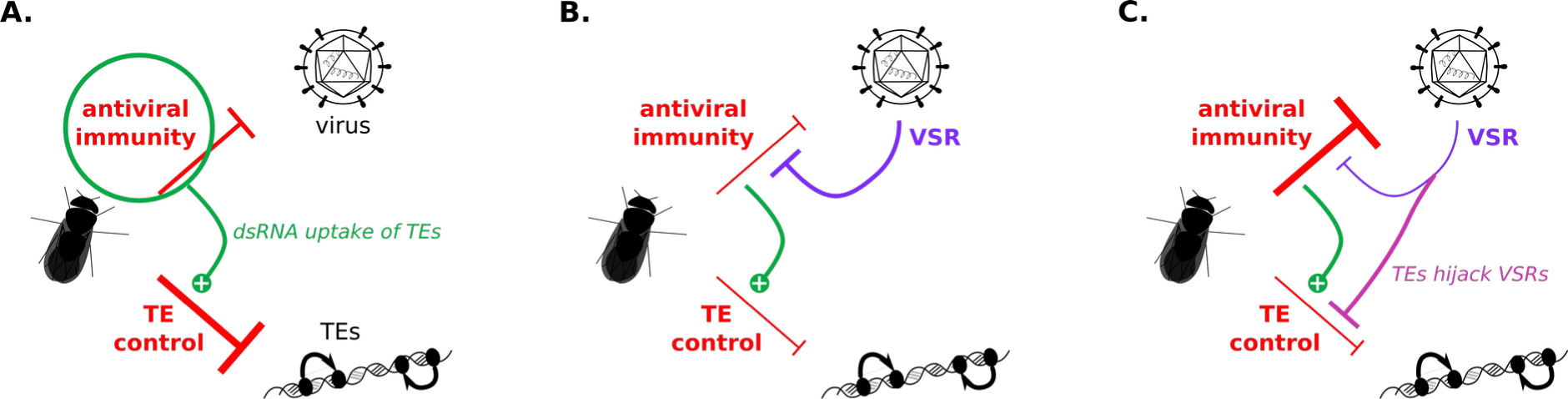
Reciprocal impacts of TE control and antiviral immunity. **A.** The fly host fights against viruses and TEs. Using SINV, we recently demonstrated that antiviral immunity enhanced TE control *via* the dsRNA uptake pathway. **B.** However, some viruses encode VSRs that inhibit antiviral immunity, such as the 1A protein produced by DCV. **C.** Here we propose that TEs hijack VSRs, which allows the release of TE control.

### TE upregulation leads to reduced DCV titers

On the other side, we wondered whether TEs could modify DCV titers. Such a study is rather difficult because it requires to make TE transcript amounts vary within the same genetic background. We thus used a sophisticated transgenic *D. melanogaster* strain, which displays an inducible *piwi* knock-down in the somatic cells surrounding the ovary (30–32) [constitutive *piwi* knock-down leads to fly sterility and thus prevents from obtaining progeny (33, 34)]. The repetitive, punctual knock-down of *piwi* along 73 successive generations – hereafter named S73, S for shift in temperature causing *piwi* knock-down– led to the accumulation of TE copies from the ZAM and gtwin families (31, 32). It was accompanied by an increase in transcript amounts for these TE families, as compared to G0-F100 –no temperature shift corresponding to the original strain– whereas gene expression remained largely unaffected (Fig S13). It has to be noted that S73 is not an isogenic strain. Instead, flies are raised as large cohorts along generations, ensuring the lowest genetic drift, and resulting in pools of flies displaying distinct, low frequency TE insertions within the same genetic background. Upon intra-thoracic injection, we found that DCV replicated earlier but at strongly reduced levels in S73 compared to G0-F100 (DCV_relative_quantity ∼ dpi + infection_status + strain; mean strain difference = 0.23; strain effect p-value = 0.004) (Fig. 4A). Fifteen days after injection, mean DCV RNA levels were 8.1-fold higher in G0-F100. Fly death rates were higher in S73 at the beginning of the infection, but the curves inverted around day 15 (log-rank test, p-value = 0.887) (Fig. 4B). Using RT-qPCR, we followed ZAM and gtwin somatic transcript amounts in both strains and both conditions (Fig. 4C and 4D). Similar to the global increase in TE transcript amounts observed in *D. melanogaster w^1118^*, we could detect that ZAM and gtwin transcript amounts in S73 are significantly higher upon DCV infection compared to mock samples around the peak of DCV titer.

**Figure 4.**
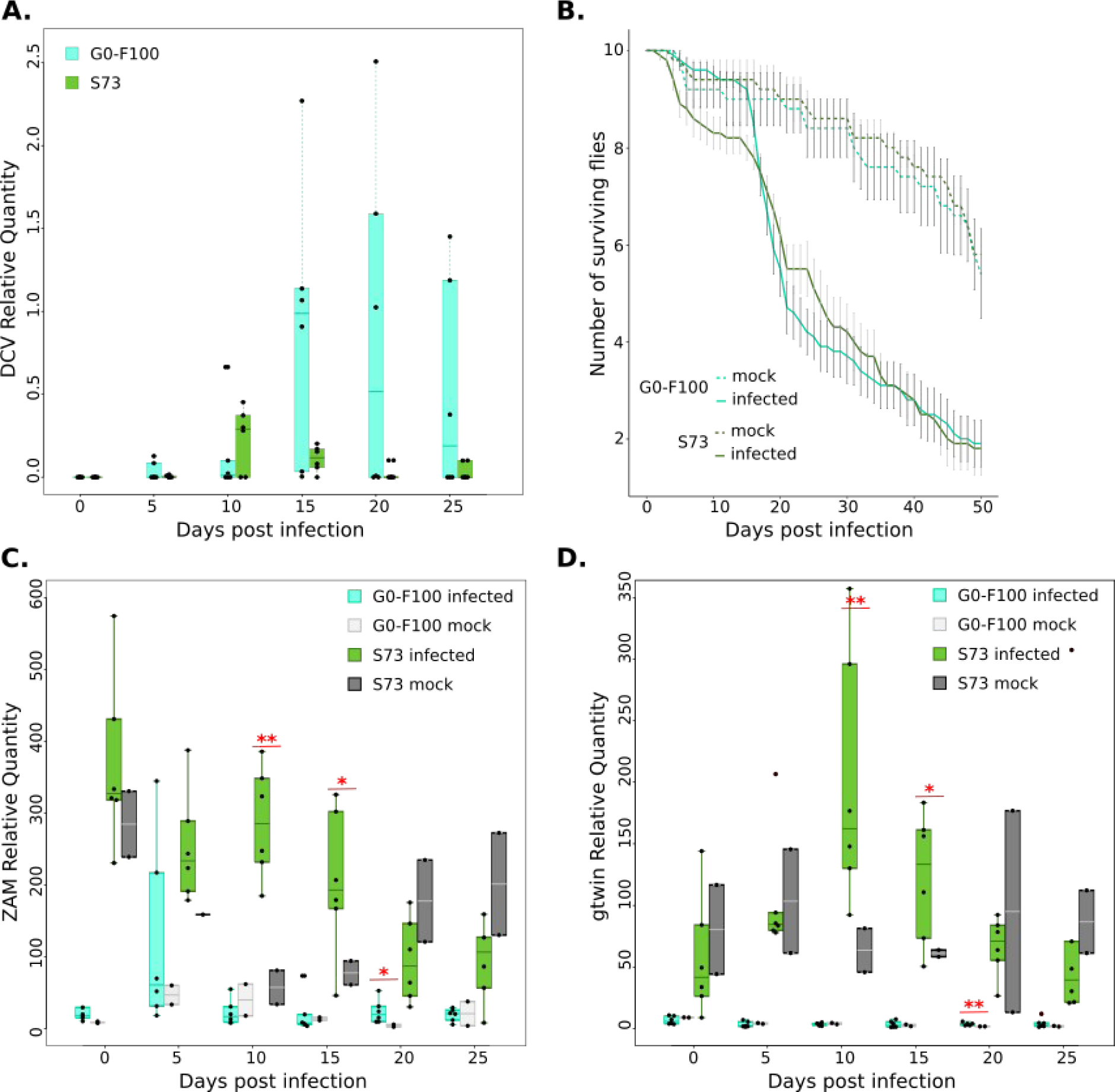
Response to DCV infection in S73 and G0-100. **A.** DCV titer in G0-F100 and S73 measured using RT-qPCR. **B.** Fly survival upon DCV and mock infections in G0-F100 and S73. **C.** ZAM transcript quantification upon DCV and mock infection in G0-F100 and S73. Stars indicate significant differences between infected and mock conditions using t-tests (p-value: 0.05 * 0.01 **). **D.** gtwin transcript quantification upon DCV and mock infection in G0-F100 and S73. Stars indicate significant differences between infected and mock conditions using t-tests (p-value: 0.05 * 0.01 **).

We propose that the higher TE transcript amounts in S73 produce increased amounts of dsRNA molecules, which can titrate 1A. The reduction in 1A availability removes DCV protection against the fly RNAi machinery; this allows the host to produce more siRNAs against the virus and therefore set up a more efficient RNAi response. This ends up in reduced viral loads (Fig. 3).

Altogether, our results suggest that a higher TE load could be beneficial in case of viral infection. At a larger evolutionary scale, such results may shed a new light on *Aedes* mosquitoes, which are known as major vectors of arboviruses –which cause no harm to them–, and are described to carry a large TE load (*ca* half of their genome). Finally, our results uncover TEs as major players of a complex host-pathogen interaction built along long-lasting coevolution.

## Material and Methods

### Drosophila strains and rearing conditions

*D. simulans* Makindu strain was previously described by Akkouche *et al.* (35) and Fablet *et al.* (36). G0-F100 and S73 transgenic *D. melanogaster* flies were previously described by Barckmann *et al.* (30) and Mohamed *et al.* (31). All experiments were performed using 3-6 day-old mated females.

Flies were reared on corn medium and maintained under standard laboratory conditions: 12/12 L/D cycle, 25°C (or 18°C for G0-F100 and S73, in order not to induce *piwi* knock-down) and 70% RH. Chronic viral infections were eliminated by bleaching the eggs, as described previously (37), except for G0-F100 and S73. Three to 6-hour-old eggs were incubated in 50% household bleach (2.6% active chlorine) for 10 minutes, washed three times for 5 minutes in deionised water and then transferred to fresh medium for adult emergence. As expected after the treatment, we could not detect amplification corresponding to the viruses using RT-PCR (SuperScript™ IV VILO™ Master Mix (without ezDNase enzyme treatment) kit and DreamTaq DNA polymerase) and the following primers: *Drosophila* A virus (DAV): 5′- AGGAGTTGGTGAGGACAGCCCA -3′ and 5′- AGACCTCAGTTGGCAGTTCGCC -3′, Nora Virus (NV): 5′- ATGGCGCCAGTTAGTGCAGACCT -3′ and 5′- CCTGTTGTTCCAGTTGGGTTCGA -3′, *Drosophila melanogaster* Sigma Virus (DmelSV): 5’- ATGTAACTCGGGTGTGACAG -3’ and 5’- CCTTCGTTCATCCTCCTGAG -3) (37). *rp49* was used as control (5′- CGGATCGATATGCTAAGCTGT -3′ and 5′- GCGCTTGTTCGATCCGTA -3′) (38). To eliminate the *Wolbachia* endosymbiotic bacteria, 3- to 6-hour-old eggs were collected and placed on fresh standard medium containing 0.25 mg/mL tetracycline hypochloride (Sigma-Aldrich), for all *Drosophila* strains. The treatment was performed during two generations and then three generations recovered on standard medium without treatment. We validated the absence of amplification by PCR using *Wolbachia 16S* primers (5′- TTGTAGCCTGCTATGGTATAACT -3′ and 5′- GAATAGGTATGATTTTCATGT -3′) (39), *Wolbachia wsp* (5′- TGGTCCAATAAGTGATGAAGAAAC -3′ and 5′- AAAAATTAAACGCTACTCCA -3′) (40) and *FtsZ* (5’- CGAGATGGGCAAAGCGATGA -3’ and 5’- ATTCCTTGCGCACCTTTCAT -3’ (41)).

### Virus production and quantification

DCV was produced and titrated in Schneider’s *Drosophila* Line DL2 cells, both kindly provided by Luis Teixeira/Ewa Chrostek. DL2 cells were kept in Schneider’s *Drosophila* Medium supplemented with 10% Fetal Bovine Serum, 1% penicillin-streptomycin 10,000 U/mL (all Gibco). Seven days after infection (MOI: 2) in Schneider’s *Drosophila* Medium supplemented with 1% penicillin-streptomycin 10,000 U/mL, the cell culture was collected and frozen at −80°C for 40 min. The culture was thawed, frozen again at −80°C for 40 min, and thawed to disrupt cells, and centrifuged at 4000 g for 10 min to remove cell debris. The supernatant was aliquoted and stored at −80°C. DCV titer was calculated by the Reed and Muench end-point calculation method: DL2 confluent cells in 96-well plates were infected with a serial 10-fold dilutions of virus suspension. The presence of active DCV was scored by cell death or clear cytopathic effects, resulting in a DCV titer of 4.22×10^9^ TCID50/mL. Similar dilutions with extracts of DL2 cells that were not inoculated with DCV did not cause any cytopathic effect in culture (42).

### Virus inoculation and survival assays

CO2-anesthetized flies were pricked in the left pleural suture on the thorax with a 0.15 mm diameter anodized steel needle dipped into the DCV solution, as described in Martinez *et al.* (26). Flies were pricked with the ‘mock’ solution –extract from uninfected DL2 cells– to control for the absence of mortality in absence of DCV infection. For the survival assays, 100 infected females and 50 ‘mock’ females were kept in rich medium vials, 10 flies per vial. Flies were transferred to fresh media every 3-4 days. The number of dead flies was recorded each day after infection.

It has to be noted that infection experiments were performed at 25°C for *D. simulans* Makindu and *D. melanogaster w^1118^* but at 18°C for *D. melanogaster* G0-F100 and S73 in order not to induce *piwi* knock-down (30). This difference in temperature leads to differences in DCV replication kinetics and in lifespan across these experiments. Accordingly, each infected sample should be analyzed in comparison with the corresponding mock sample obtained in the exact same experimental conditions.

### RNA extraction and RT-qPCR

To quantify DCV and TE expression, total RNAs were extracted individually for 6 infected and 2 mock flies at 0, 2, 4, 6 and 8 dpi using Qiazol and the RNeasy mini kit (Qiagen). Purified RNAs were processed using Turbo DNase (Ambion DNAfree kit). Reverse transcription was performed on 4 μL of total RNAs using SuperScript IV VILO Master Mix (Invitrogen). qPCR was performed using 2 μL of cDNA, 0.5 μL of each primer (DCV : 5’- GACACTGCCTTTGATTAG -3’ and 5’- CCCTCTGGGAACTAAATG -3’, ZAM : 5’- CTACGAAATGGCAAGATTAATTCCACTTCC -3’ and 5’- CCCGTTTCCTTTATGTCGCAGTAGCT -3’, gtwin : 5’- TTCGCACAAGCGATGATAAG -3’ and 5’- GATTGTTGTACGGCGACCTT -3’) and 5 μL of SsoADV SYBR® Green Supermix (Bio Rad) in a QuantStudio™ 6 Flex Real-Time PCR System (Applied Biosystems™) and following : 30 sec at 95 °C; 40 cycles of 95 °C for 15 sec and 58 °C for 30 sec. For standardization, we tested the *rp49*, *Adh*, and *EF1* genes, and kept *rp49* for further experiments because it displayed the highest stability across all conditions.

### RNA-seq

Thirty pairs of ovaries were carefully and manually dissected and separated from the rest of the bodies (‘carcasses’) at 4 dpi. Total RNAs from 30 pairs of ovaries or 30 carcasses were extracted according to the protocol described above, using Qiazol, the RNeasy Mini Kit (Qiagen) and TurboDNAse (Ambion DNAfree kit) on the total amount of eluate to eliminate residual DNA. RNA quality was validated using a Bioanalyzer (Agilent). Library preparation was performed at the GenomEast platform at the Institute of Genetics and Molecular and Cellular Biology (IGBMC, France) using Illumina Stranded mRNA Prep Ligation - Reference Guide - PN 1000000124518. Libraries were sequenced as paired-end 100 base reads (HiSeq 4000: Illumina).

For S73 and G0-100 samples, total RNA was extracted from 3-16 h embryos using TRIzol. 4 mg of total RNA was subjected to Ribo-Zero ribosomal depletion. RNA was further purified using RNA Clean & Concentrator-5. Libraries were prepared using the Illumina Stranded mRNA Prep Ligation kit and 50 nt paired-end read sequencing were performed by MGX sequencing services (Montpellier, France) on a SP flow cell using Novaseq 6000. The RNA-seq experiments were performed on three biological replicates.

Read alignment was performed using Hisat2 (43) on *D. melanogaster* and *D. simulans* reference genes available from FlyBase, versions r6.16 and r2.02, respectively, and previously masked using RepeatMasker. The reference sequence used for DCV was NC_001834.1 from GenBank. Alignments were processed using SAMtools (44). Gene count tables were generated using eXpress (45). TE count tables were obtained using the TEcount module of TEtools (46) and the same TE reference files as used in (18), and corresponding to 237 TE families. Gene and TE count tables were concatenated and then analyzed using the DESeq2 R package (version 4.3) (47). Four complete count tables were produced ({*D. melanogaster* carcasses}, {*D. melanogaster* ovaries}, {*D. simulans* carcasses}, and {*D. simulans* ovaries}). Each complete table was then analyzed by the DESeq2 procedure, ensuring that TE counts are normalized the same way as gene counts. The gene ontology (GO) analysis of RNA-seq data was performed on the differentially expressed genes with the gseGO function of the clusterProfiler package (48, 49). The GO terms were obtained using the R package org.Dm.eg.db v3.10.0 as database (50), corresponding to all *D. melanogaster* annotated genes. We converted *D. simulans* genes into their *D. melanogaster* orthologs using the orthology file from FlyBase ftp://ftp.flybase.net/releases/FB2020_04/precomputed_files/orthologs/dmel_orthologs_in_drosophila_species_fb_2020_04.tsv.gz. The GO overrepresentation analysis of biological process (BP) was performed with adjusted p-values by the FDR method, p-value cutoff at 0.05, and a minimum of 3 genes per term.

### small RNA-seq

Ovaries of each female fly were carefully and manually dissected at 4 dpi. Small RNAs from 30 pairs of ovaries or 30 carcasses were extracted using the TraPR (Trans-kingdom, rapid, affordable Purification of RISCs) small RNA Isolation kit (Lexogen), as described previously (51). The TraPR method allows the specific isolation of fully functional and physiologically relevant interfering RNAs (microRNAs, piRNAs, siRNAs and scnRNAs), by anion exchange chromatography. Size selection (18-40 nt) was performed on gel at the sequencing platform. Purified small RNAs were used for library preparation, using TruSeq Small RNA Sample Preparation Guide - PN 15004197. Libraries were sequenced on an Illumina HiSeq 4000 sequencer as single-end 50 base reads.

Sequencing adapters were removed using cutadapt (52) -a TGGAATTCTCGGGTGCCAAGGAACTCCAGTCACTTA -m 1. Size filtering was performed using PRINTSEQ (53), in order to distinguish 21 nt-long reads (considered as siRNAs) from 23-30 nt-long reads (considered as piRNAs).

TE count tables for sense and antisense alignments were obtained using a modified version of the TEcount module (36) of TEtools (46) on the same TE reference sequences as above. Ping-pong signatures were checked using signature.py with min_size = 23 and max_size = 30 options (54). Read count normalization using microRNA counts as supposed invariants is a common strategy. However, it has been shown that some microRNA amounts could be affected by DCV infection (55). This is why we performed three different normalization strategies : 1) normalization using all microRNA sequences annotated from FlyBase (19-39 nt-long reads mapped using bowtie --best on FlyBase reference sequences dmel-all-miRNA-r6.16.fasta and dsim-all-miRNA-r2.02.fasta), 2) normalization using microRNA sequences but excluding those described to vary upon DCV infection in Monsanto-Hearnes *et al.* (55), 3) normalization using the endo-siRNAs described by Malone *et al.* (56), as we previously did in Roy *et al.* (18)). However, this last normalization procedure is only possible in *D. melanogaster* because endo-siRNA producing loci are not annotated in *D. simulans*. All normalization strategies provided similar patterns (Fig S8-S10).

## Data availability

The RNA-seq and small RNA-seq data generated in this study have been submitted to the NCBI BioProject database under accession number PRJNA996035.

## Acknowledgments

This work was performed using the computing facilities of Laboratoire de Biométrie Biologie Evolutive / Pôle Rhône-Alpes de bioinformatique (CC LBBE/PRABI) and the Symbiotron Pasteur platform, Laboratoire Biologie Fonctionnelle Insectes et Interactions (BF2i, Lyon). Sequencing was performed by the GenomEast platform, a member of the ‘France Génomique’ consortium (ANR-10-INBS-0009). We thank Matthieu Boulesteix, Miriam Merenciano, Cristina Vieira for helpful discussions. We thank Nelly Burlet, Sonia Janillon, Corinne Régis, Zaïnab Belgaidi, Carole Monégat, and Agnès Vallier for technical help. We thank also Clément Gilbert, Anna Zaidman-Rémy, and Feth el Zahar Haichar for helpful discussions. This work was funded by Agence Nationale de la Recherche (TEMIT grant) to M.F. and by the Agence Nationale de la Recherche (ANR-19-CE12-0012 -Top53 grant) to S.C.. M.V. was funded by CNRS-Japon PhD Joint Program.

## Notes

### Competing Interest Statement

The authors have declared no competing interest.

