## Supplementary Material for "Intricate interactions between antiviral immunity and transposable element control in *Drosophila*"

### Supplementary Information

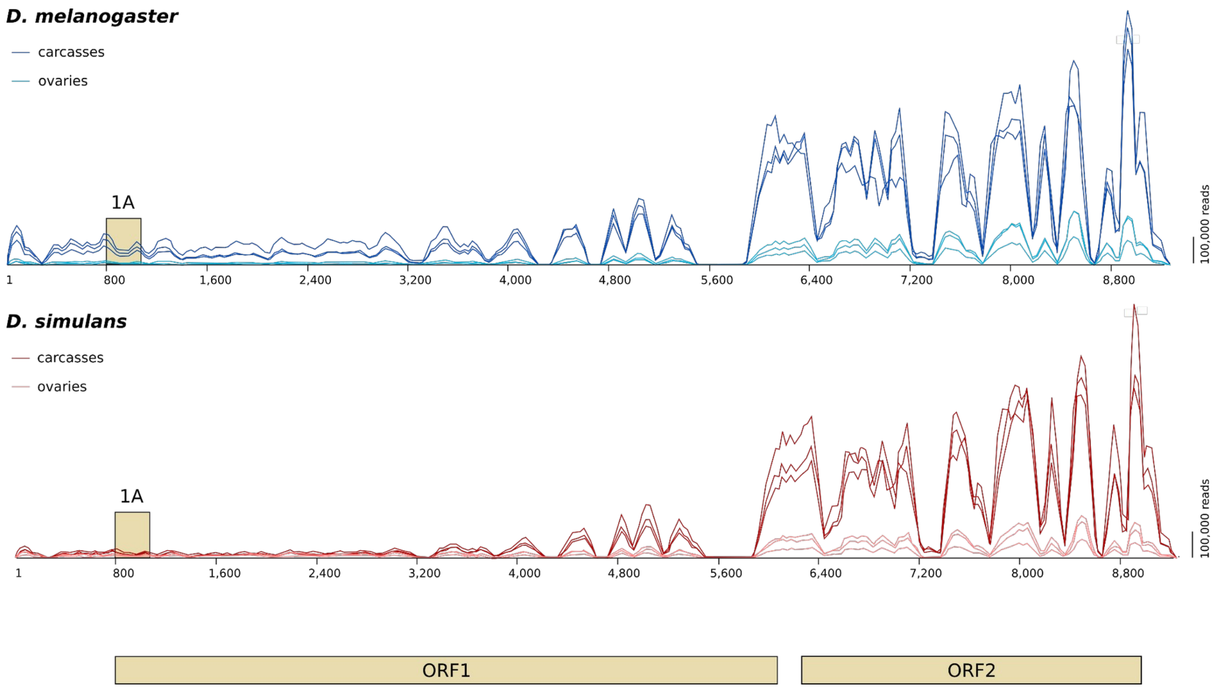


**Figure S1. RNA-seq reads mapping along the DCV genome.** The three biological replicates are shown for each condition. The annotation of the reference DCV sequence (NCBI NC_001834.1) is provided in beige at the bottom, to scale. ORF1 encodes non structural proteins; ORF2 encodes structural proteins. The box corresponding to the 5’ end of ORF1 shows the position of 1A VSR.

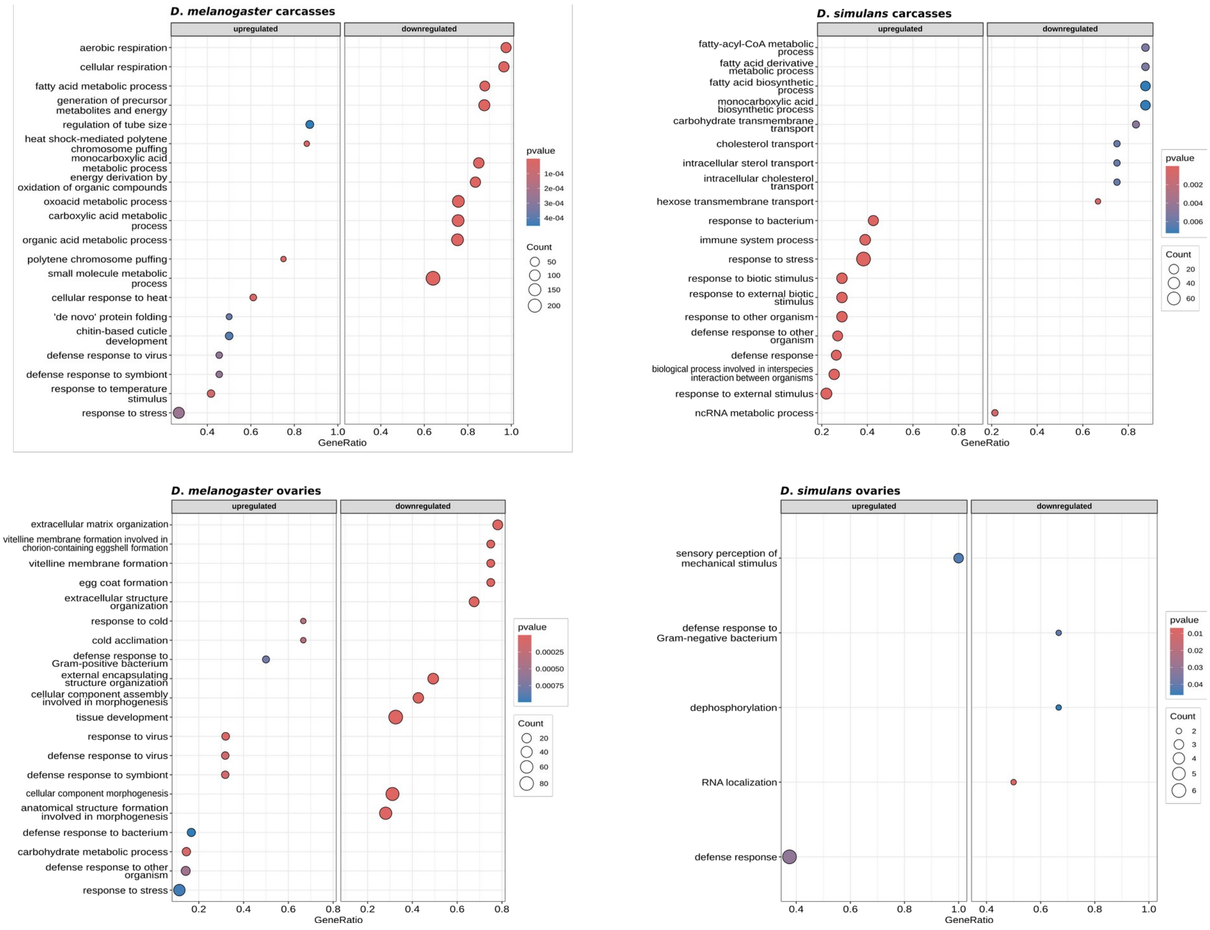


**Figure S2. GO term analysis upon infection in RNA-seq samples.** Fifteen most enriched GO terms in *D. melanogaster* carcasses (upper left panel) and *D. simulans* carcasses (upper right panel) upon DCV infection. Fifteen most enriched GO terms in *D. melanogaster* ovaries (lower left panel) and *D. simulans* ovaries (lower right panel) upon DCV infection. GeneRatio is the number of DE genes reported to the total number of genes in the corresponding GO term.

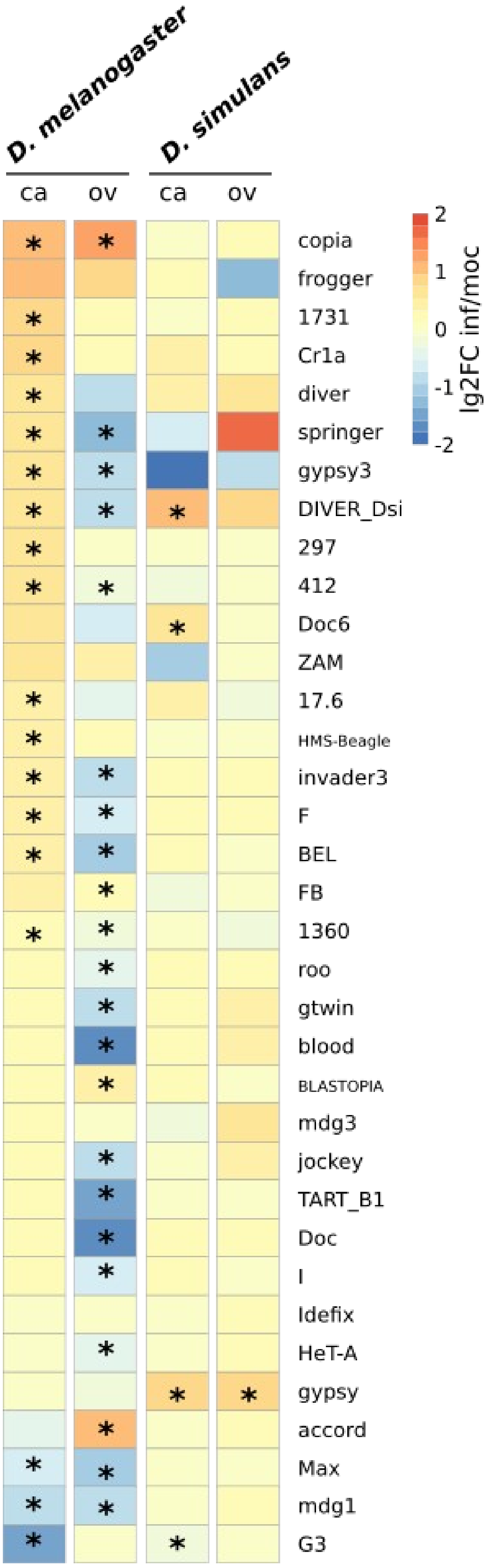


**Figure S3. Heatmap of log2FC values between infected and mock conditions for some TE families.** Stars indicate TE families that are significantly differentially expressed (DESeq2 adjusted p-value $< 0.05$).

**Table S4. Contribution of TE classes to differentially expressed TE families.**

**A.** Median values across TE families of log2FC in read counts between infected and mock conditions, from RNA-seq data. P-values correspond to the paired Wilcoxon test comparing read counts in infected *vs* mock conditions across TE families.

| | LTR | | LINE | | DNA | |
|---|---|---|---|---|---|---|
| Condition | median log2FC | p | median log2FC | p | median log2FC | p |
| *D. melanogaster* carcasses | 0.292 | 3.1e-6 | 0.230 | 9.1e-5 | 0.306 | 9.1e-4 |
| *D. melanogaster* ovaries | -0.349 | 7.2e-5 | -0.477 | 2.7e-4 | -0.240 | 0.002 |
| *D. simulans* carcasses | 0.044 | 0.052 | 0.059 | 0.260 | 0.054 | 0.691 |
| *D. simulans* ovaries | 0.152 | 7.8e-8 | 0.131 | 1.7e-4 | -0.046 | 0.446 |

**B.** Distributions of the numbers of differentially expressed (adjusted p-value < 0.05) TE families according to TE class, and Fisher exact tests comparing distributions across classes.

| | | LTR | LINE | DNA | p-value |
|---|---|---|---|---|---|
| *D. melanogaster* carcasses | log2FC > 0 | 31 | 7 | 5 | 1 |
| | log2FC < 0 | 6 | 1 | 0 | |
| *D. melanogaster* ovaries | log2FC > 0 | 7 | 2 | 0 | 0.619 |
| | log2FC < 0 | 24 | 13 | 4 | |
| *D. simulans* carcasses | log2FC > 0 | 4 | 1 | 1 | 0.190 |
| | log2FC < 0 | 1 | 3 | 0 | |
| *D. simulans* ovaries | log2FC > 0 | 1 | 1 | 0 | 1 |
| | log2FC < 0 | 0 | 0 | 0 | |

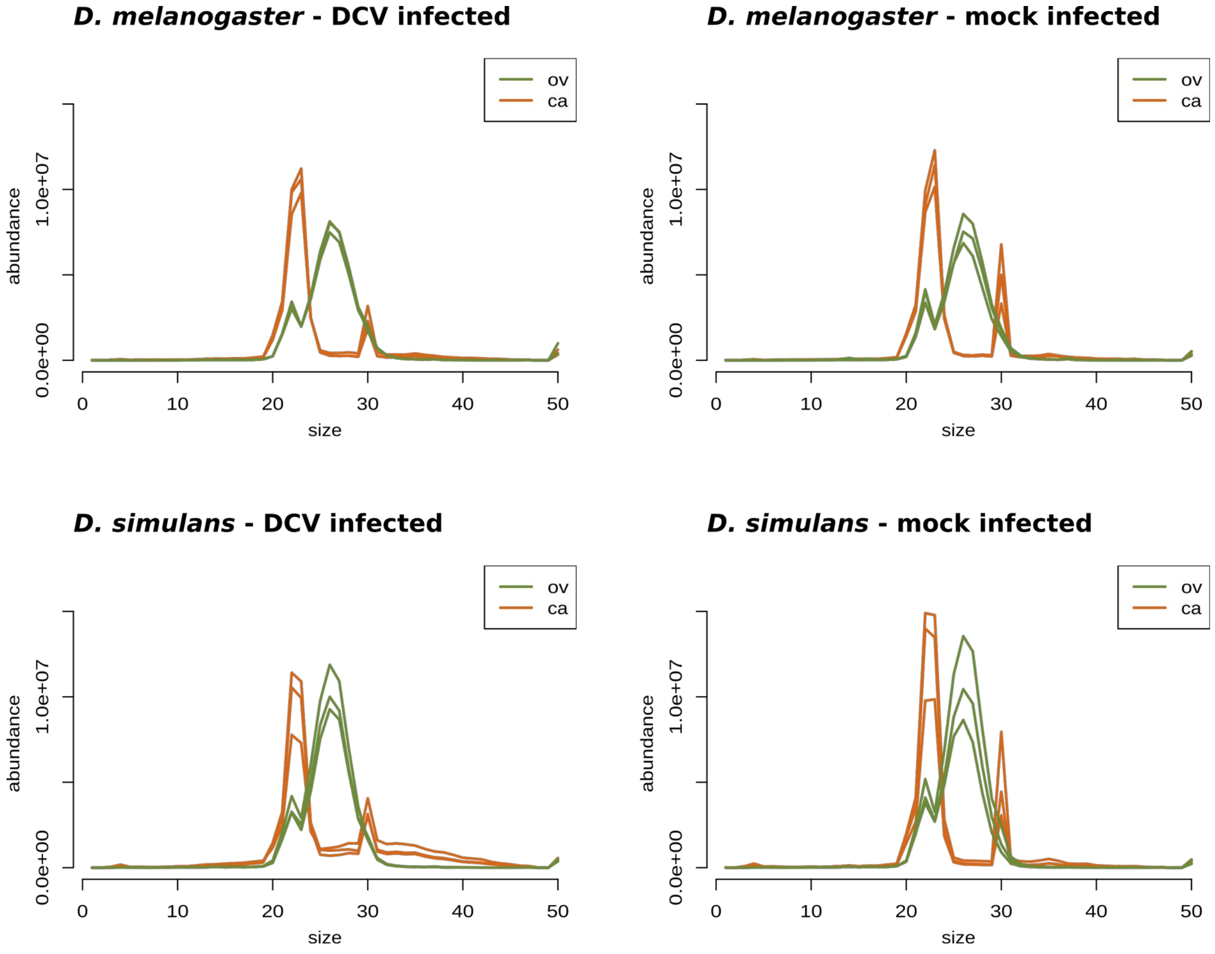


**Figure S5. Size distribution of small RNA-seq reads.** Abundance in raw read numbers. As expected, reads corresponding to carcass samples (brown) were 21 nt-long in the majority, *i.e.* the expected size for siRNAs, and reads corresponding to ovary samples (green) in the majority displayed sizes of 21 nt (siRNAs) and 23 to 30 nt (piRNAs). The 30 nt-long peaks in carcass samples corresponded to rRNAs. Three biological replicates.

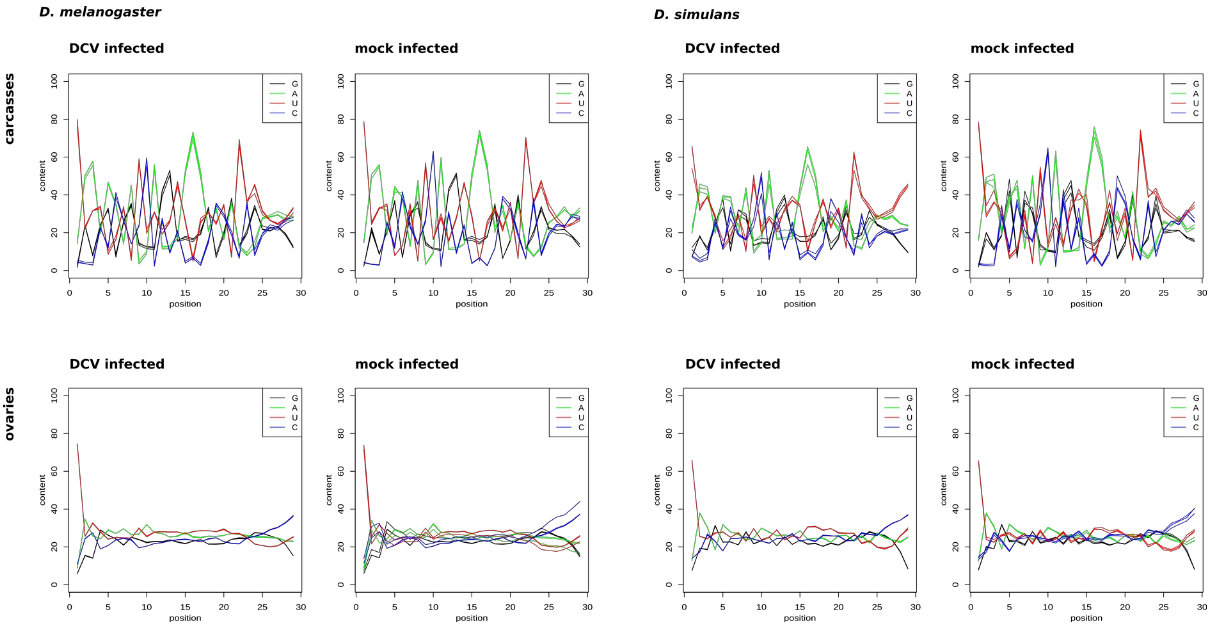


**Figure S6. Per base nucleotide composition of 23 to 29 nt-long small RNA-seq reads.** All samples displayed an enrichment in U at the first position, as expected for functional piRNAs.

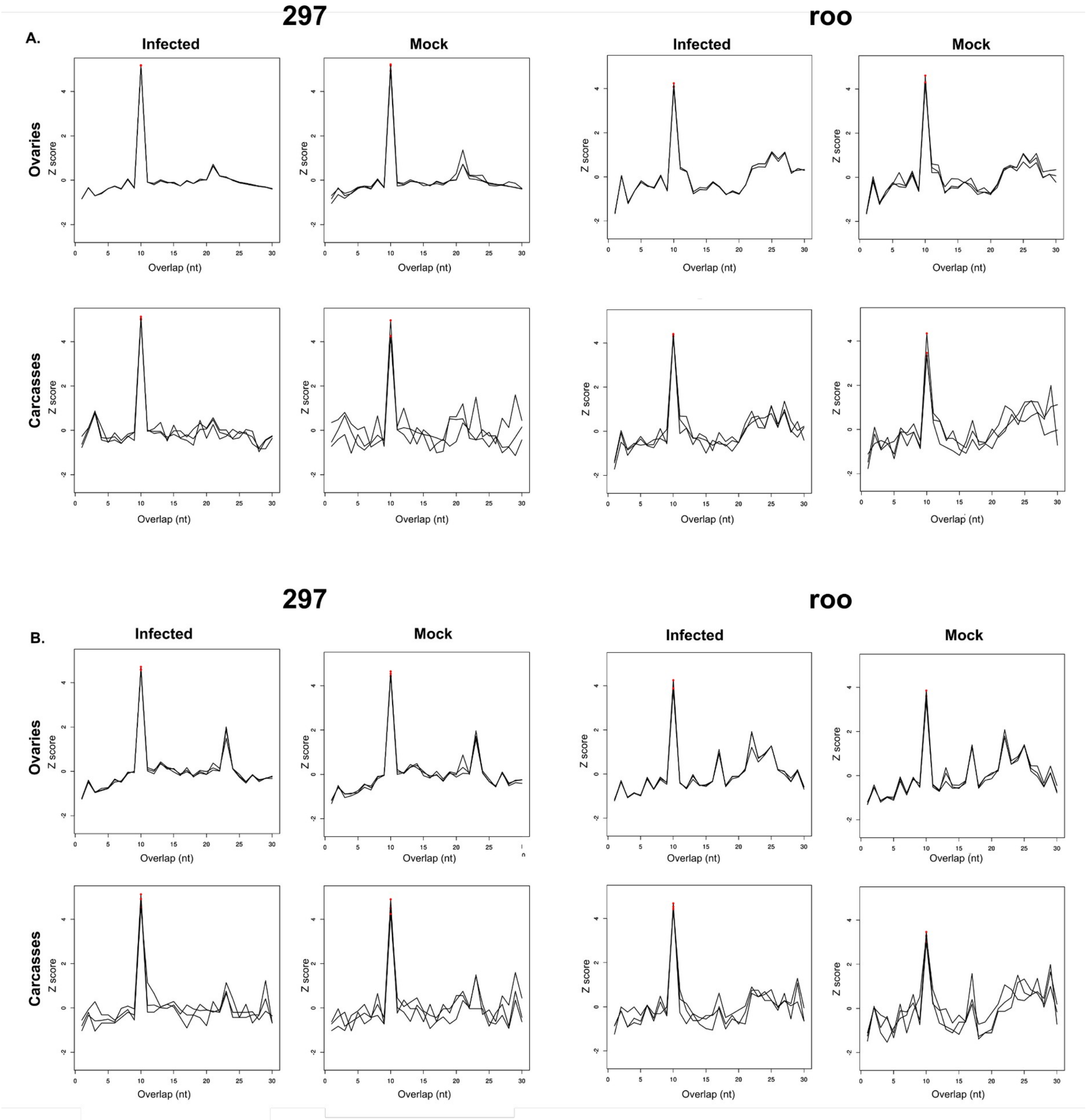


**Figure S7. Ping-pong signatures for the roo and 297 TE families in *D. melanogaster* (A) and *D. simulans* (B)**. We looked for ping-pong signatures using signature.py with the options min_size = 23 and max_size = 30 (54). Alignments of 23-30 nt reads were performed using bowtie --best on 297 and roo consensus sequences available at https://github.com/bergmanlab/drosophila-transposons/blob/master/releases/D_mel_transposon_sequence_set_v10.2.fa. X-axes show the length of the overlaps and Y-axes show the corresponding z-scores, *i.e.* the standardized numbers of pairs found for each length of overlap (($p_i$ – mean(p)) / standard_deviation(p), where $p_i$ is the number of pairs of i nt overlaps).

The peak at overlap 10 nt is indicative of an efficient ping-pong loop. Three replicates are shown.

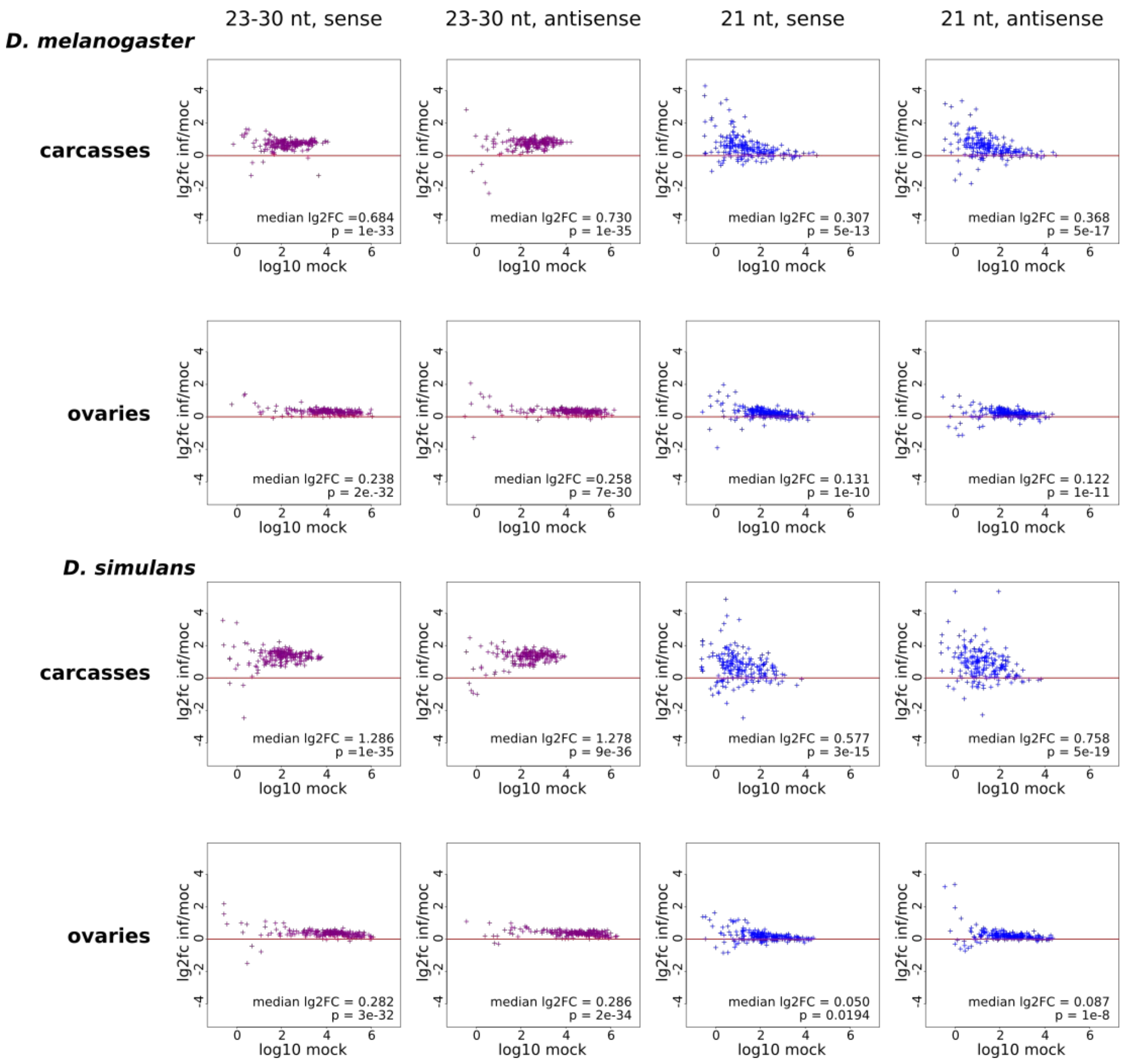


**Figure S8. TE-derived small RNAs log2FC between infected and mock conditions**. Normalization using the complete set of microRNA sequences. Paired Wilcoxon tests p-values comparing infected and mock counts.

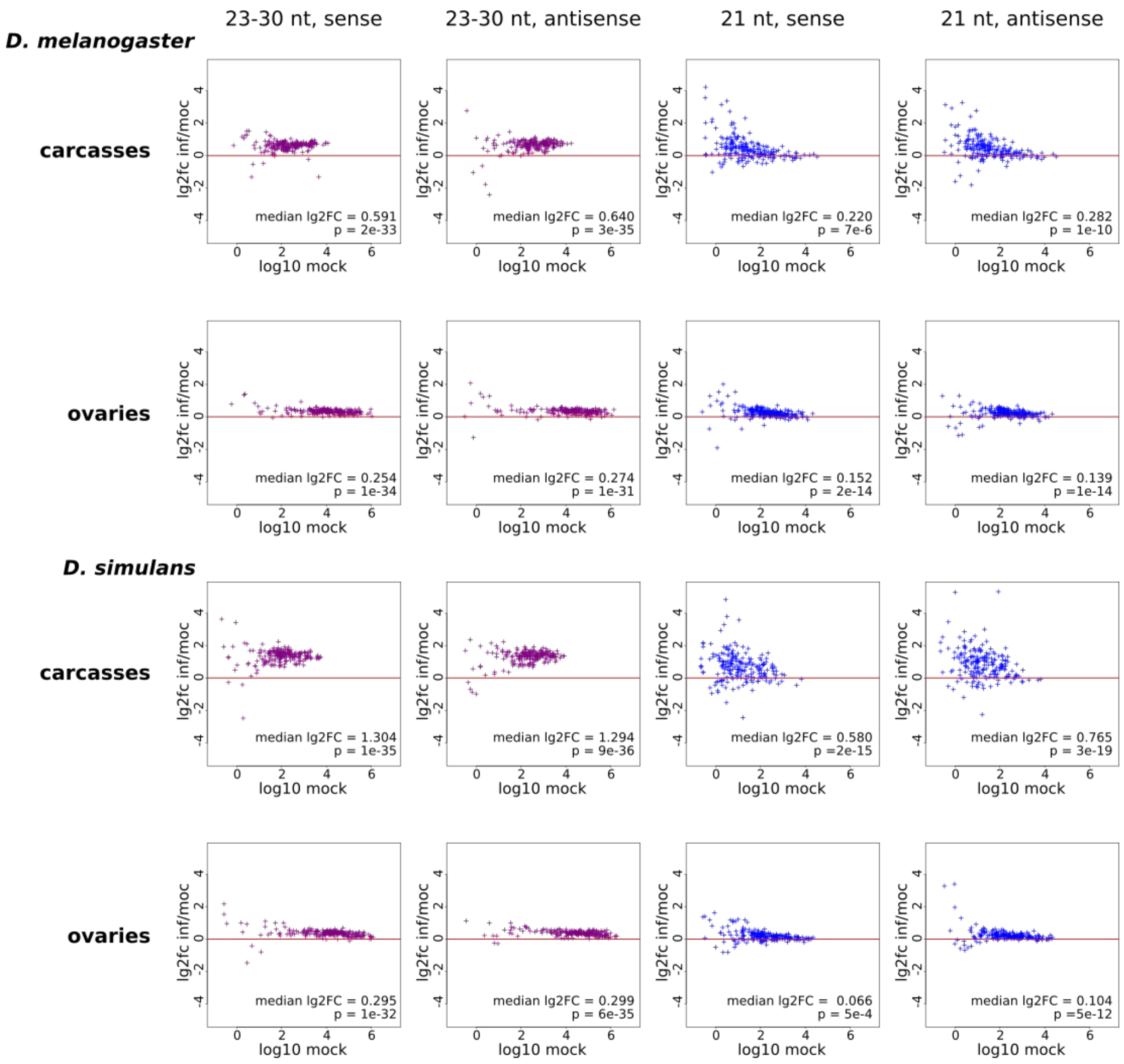


**Figure S9. TE-derived small RNAs log2FC between infected and mock conditions**. Normalization using microRNA sequences and removing the microRNAs identified in (55) as modified upon DCV infection. Paired Wilcoxon tests p-values comparing infected and mock counts.

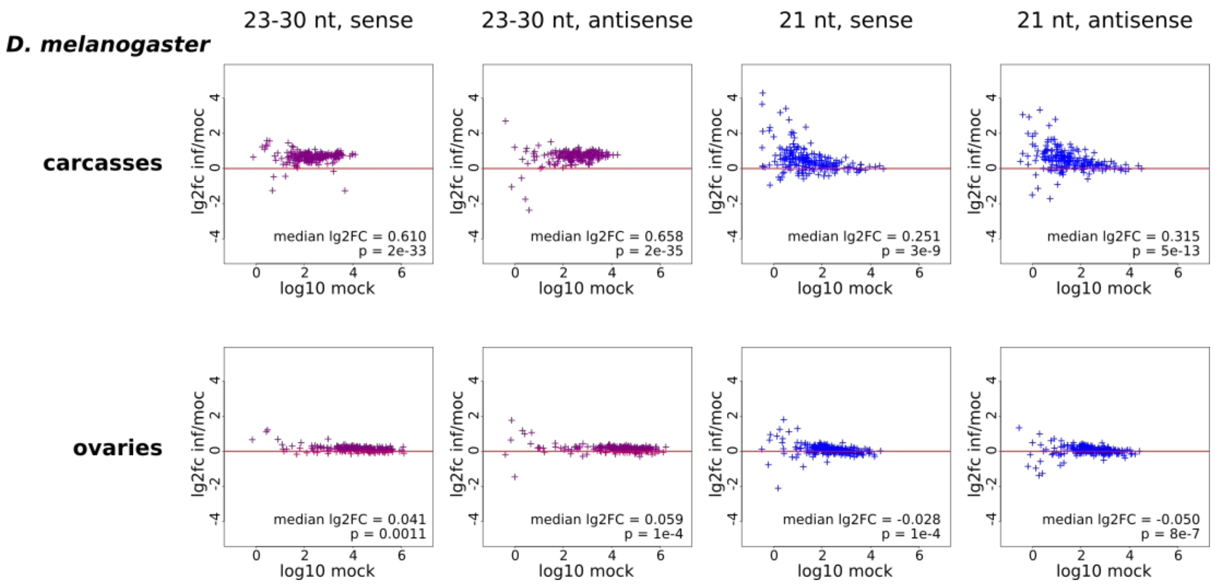


**Figure S10. TE-derived small RNAs log2 Fold Changes between infected and mock conditions**. Normalization using endo-siRNAs, as defined by (56). This normalization can only be performed on *D. melanogaster* samples because endo-siRNA clusters are not annotated for *D. simulans*. Paired Wilcoxon tests p-values comparing infected and mock counts.

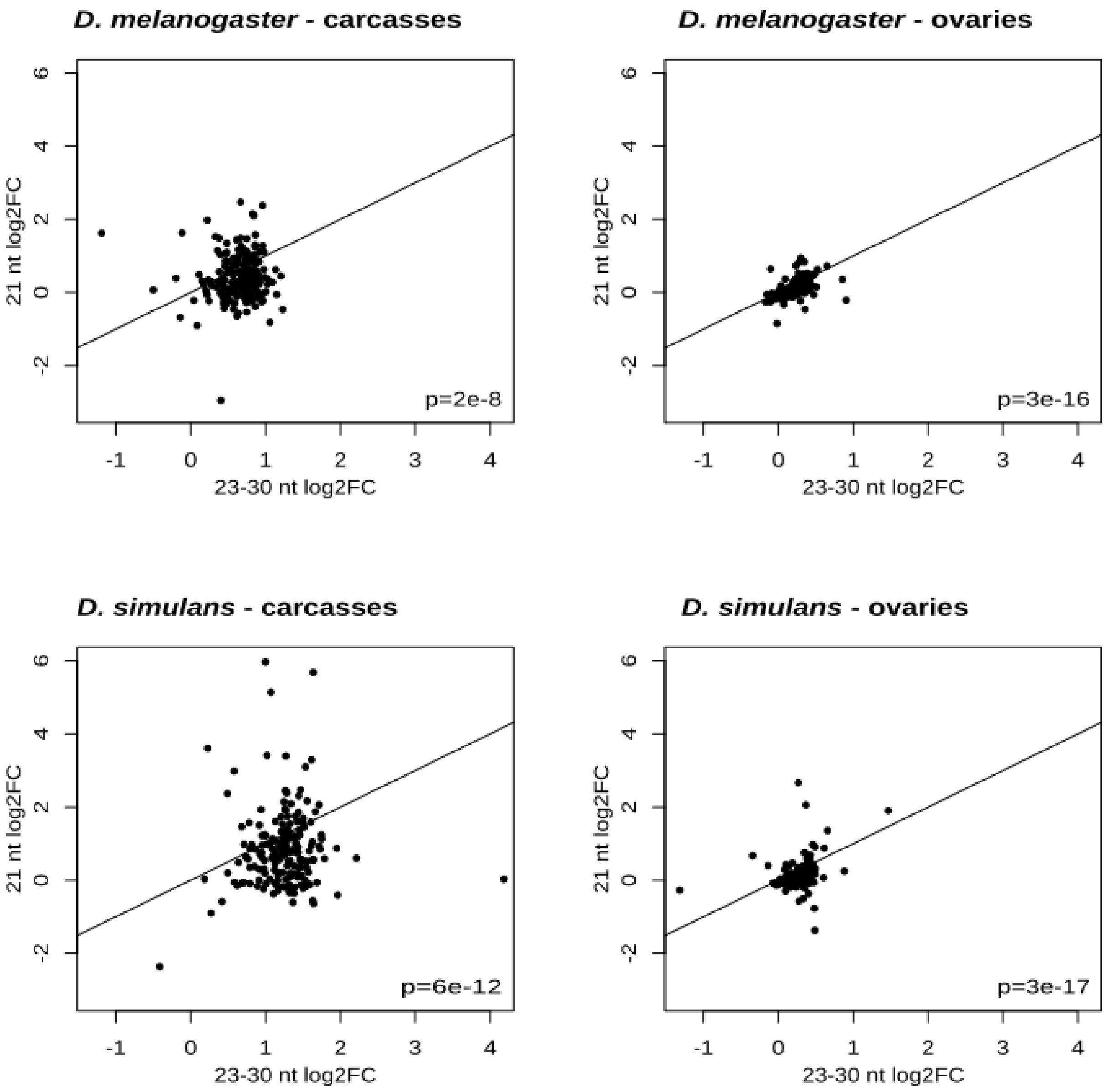


**Figure S11. Comparison of log2 Fold Changes between TE-derived siRNAs and piRNAs.** In all conditions, log2 Fold Changes were higher for TE-derived 23-30 nt long small RNAs (piRNAs) compared to TE-derived 21 nt long small RNAs (siRNAs). Paired Wilcoxon test p-values comparing 21 nt log2FC and 23-30 nt log2FC.

**Table S12. Dependency tests between TE transcript log2FC and TE-derived small RNA log2FC**.

*D. melanogaster* carcasses

| si sense | | | | si anti | | | |
|---|---|---|---|---|---|---|---|
| | sRNA lg2fc <0 | sRNA lg2fc >0 | p | | sRNA lg2fc <0 | sRNA lg2fc >0 | p |
| RNA lg2fc <0 | 10 | 43 | 0.829 | RNA lg2fc <0 | 10 | 41 | 0.959 |
| RNA lg2fc >0 | 30 | 109 | | RNA lg2fc >0 | 30 | 111 | |
| **pi sense** | | | | **pi anti** | | | |
| | sRNA lg2fc <0 | sRNA lg2fc >0 | p | | sRNA lg2fc <0 | sRNA lg2fc >0 | p |
| RNA lg2fc <0 | 1 | 52 | 1 | RNA lg2fc <0 | 3 | 51 | 0.029 |
| RNA lg2fc >0 | 3 | 138 | | RNA lg2fc >0 | 0 | 141 | |

*D. melanogaster* ovaries

| si sense | | | | si anti | | | |
|---|---|---|---|---|---|---|---|
| | sRNA lg2fc <0 | sRNA lg2fc >0 | p | | sRNA lg2fc <0 | sRNA lg2fc >0 | p |
| RNA lg2fc <0 | 35 | 107 | 0.843 | RNA lg2fc <0 | 36 | 107 | 0.561 |
| RNA lg2fc >0 | 15 | 40 | | RNA lg2fc >0 | 11 | 44 | |
| **pi sense** | | | | **pi anti** | | | |
| | sRNA lg2fc <0 | sRNA lg2fc >0 | p | | sRNA lg2fc <0 | sRNA lg2fc >0 | p |
| RNA lg2fc <0 | 5 | 138 | 0.822 | RNA lg2fc <0 | 10 | 133 | 0.867 |
| RNA lg2fc >0 | 3 | 52 | | RNA lg2fc >0 | 5 | 51 | |

*D. simulans* carcasses

| si sense | | | | si anti | | | |
|---|---|---|---|---|---|---|---|
| | sRNA lg2fc <0 | sRNA lg2fc >0 | p | | sRNA lg2fc <0 | sRNA lg2fc >0 | p |
| RNA lg2fc <0 | 18 | 64 | 0.799 | RNA lg2fc <0 | 18 | 62 | 0.244 |
| RNA lg2fc >0 | 28 | 86 | | RNA lg2fc >0 | 17 | 97 | |
| **pi sense** | | | | **pi anti** | | | |
| | sRNA lg2fc <0 | sRNA lg2fc >0 | p | | sRNA lg2fc <0 | sRNA lg2fc >0 | p |
| RNA lg2fc <0 | 2 | 81 | 0.763 | RNA lg2fc <0 | 1 | 83 | 0.860 |
| RNA lg2fc >0 | 1 | 116 | | RNA lg2fc >0 | 3 | 114 | |

| *D. simulans* ovaries | si sense | | | | si anti | | | |
|---|---|---|---|---|---|---|---|---|
| | | sRNA lg2fc <0 | sRNA lg2fc >0 | p | | sRNA lg2fc <0 | sRNA lg2fc >0 | p |
| | RNA lg2fc <0 | 28 | 37 | 0.363 | RNA lg2fc <0 | 17 | 46 | 0.962 |
| | RNA lg2fc >0 | 48 | 88 | | RNA lg2fc >0 | 39 | 98 | |
| | pi sense | | | | pi anti | | | |
| | | sRNA lg2fc <0 | sRNA lg2fc >0 | p | | sRNA lg2fc <0 | sRNA lg2fc >0 | p |
| | RNA lg2fc <0 | 3 | 62 | 0.831 | RNA lg2fc <0 | 2 | 64 | 0.938 |
| | RNA lg2fc >0 | 4 | 134 | | RNA lg2fc >0 | 6 | 131 | |

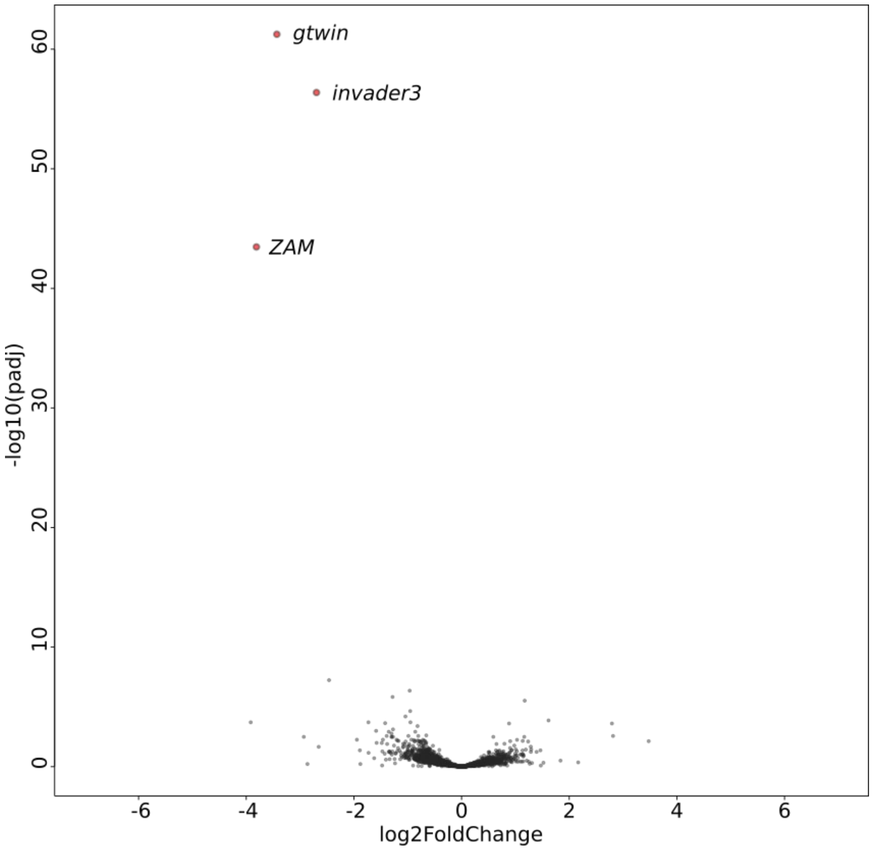


**Figure S13. Differential expression analysis of G0-100 *vs* S73, Volcano plot**. RNA-seq data obtained from embryos on biological triplicates. S73 and G0-100 transcriptomes essentially differ regarding gtwin, invader3, and ZAM TE families (over-expressed in S73), while gene and other TE family levels are conserved (gray dots).
